# Piezo ion channel activation increases the release of therapeutic extracellular vesicles after mechanical stimulation in bioreactors

**DOI:** 10.1101/2025.01.09.632205

**Authors:** André Cronemberger Andrade, Oriane Le Goas, Shony Lemieux, Alice Grangier, Alice Nicolai, Chiara Guerrera, Christopher Ribes, Estelle Suply, Jeanne Volatron, Florence Gazeau, Amanda. K.A. Silva

## Abstract

Enhancing protocols and methods for producing therapeutic extracellular vesicles (EVs) in bioreactors is crucial to achieve scalable production while ensuring both quality and quantity. Studies have shown that mechanical stress can promote EV release, although the underlying mechanisms remain largely unclear. Here we investigated which mechanisms are responsible for the increase of EV production under shear stress. EVs were produced from adipose tissue-derived stromal cells (ASCs), also described as mesenchymal stromal cells (MSCs) that can support the regeneration of injured tissues via several paracrine factors. The cultures were treated with GsMTx4, an inhibitor which blocks the Piezo1 ion channels, or with YODA1, its agonist, to assess if mechanical or shear forces pathways are involved in enhancing the EV release. EVs were quantified and characterized after high shear (HS) stimulation compared with no shear stress stimulation (3D) and standard cultures methods (2D). These experiments showed that, after mechanical stimulation by shear stress, EV production increased in bioreactors and this effect was blocked by the inhibition of Piezo1 ion channels with GsMTx4 (88%) with no impact on cell viability. Consistently, the agonist YODA1 increased the EV production (149%). The implications of these findings are significant, especially for regenerative medicine and cellular therapies, where the efficient production of high-quality EVs is crucial. By understanding turbulence-induced shear stress and the natural mechanotransductive pathways within cells, it may be possible to optimize the production of therapeutic EVs, paving the way for more effective treatments in the future.

## Introduction

Extracellular vesicles (EVs) are nanosized membrane-bound particles naturally secreted by cells, playing a crucial role in intercellular communication ^1^. They carry diverse biomolecules, including proteins, lipids, and nucleic acids, which reflect their cell of origin and physiological state ^2^. EVs are involved in various biological processes, such as immune modulation, tissue repair, and tumor progression, making them pivotal in health and disease ^1,3^. Their unique ability to transfer functional molecules to target cells has garnered significant attention in the fields of regenerative medicine, drug delivery, and biomarker discovery ^4^.

One of the primary challenges in EV research and application is the efficient and scalable production of high-quality EVs ^5^. Traditional 2D cell culture systems are limited by their low production yields, making them unsuitable for clinical-grade EV manufacturing of human doses ^5^. Innovative approaches, such as 3D culture systems and bioreactors, have been explored to enhance EV production ^5^. The ability to increase EV production efficiently while maintaining their functional integrity and therapeutic potential is essential for advancing EV-based technologies.

Mesenchymal stem cells (MSCs) are a prominent source of therapeutic EVs due to their immunomodulatory and regenerative properties ^6^. Unlike their parent cells, MSC-derived EVs (MSC-EVs) offer several advantages: they are non-replicative, minimizing the risk of tumor formation; they can cross biological barriers, such as the blood-brain barrier; and they are easier to store and transport ^6^. MSC-EVs have shown promise in treating a wide range of conditions, including cardiovascular diseases, neurodegenerative disorders, and inflammatory diseases ^7–11^. They mediate their effects by modulating immune responses, promoting tissue repair, and delivering bioactive molecules to damaged or diseased tissues.

Mechanical stimulation, such as shear stress, has been shown to significantly enhance the production of MSC-EVs, surpassing the efficiency of current standard methods ^12–14^. According to studies from our group, the application of turbulence within a controlled bioreactor environment induces shear stress on MSCs, which in turn promotes the release of EVs ^15–17^. This finding highlights the potential of mechanical stimulation as a superior method for EV bioproduction, particularly in therapeutic applications.

In this study, we aimed to uncover the mechanisms driving EV release under shear stress, a process that remains poorly understood. To achieve this, we analyzed proteome cargo of the EVs to identify potential clues about the molecular pathways involved. The findings from this analysis pointed toward a mechanosensitive hypothesis, suggesting that mechanotransduction may play a key role in regulating EV secretion. Building on this hypothesis, we investigated the contribution of mechanosensitive ion channels, particularly Piezo1, to EV release. This paper presents our findings, shedding light on the interplay between shear stress and EV biogenesis.

## Results

### High shear stimulation in bioreactors increases shear-stress related protein Piezo1 in EVs and cells

To gain insight on the biogenesis mechanisms at play, the productions of EVs from human MSCs using three culture methods (culture flasks (2D), stirred-tank bioreactors (3D) and stirred-tank bioreactors with mechanical stimulation by high agitation speed (HS)) were compared and proteomic analysis was performed. In previous studies, 3D culture was shown to increase EV yield ^18,19^. Particle production was significantly higher in HS conditions compared to 2D (by factor of 10.33) and 3D (by a factor of 5,65), indicating that under shear stress cells produce more EVs (Fig. 1A). Principal component analysis (PCA) of the EVs proteome patterns revealed distinct clustering of replicates for each culture method, confirming that the protein expression profiles were method-dependent among the EVs (Fig. 1B). Where the higher variation is found in EVs derived from 2D when compared with EV derived from 3D and HS (Fig. 1B). EVs derived from 3D cultures showed less variation compared to EVs derived from HS (Fig. 1B). A Venn diagram was used to visually represent the overlap and distinctiveness of significantly expressed proteins across the three culture methods: 2D, 3D, and high-shear (HS) conditions. The Venn diagram highlighted the shared proteins among the methods, providing insight into the common molecular signatures potentially related to EV production in all conditions (Supplementary Fig. 1A). Simultaneously, it underscored the unique protein expressions specific to each method, reflecting the influence of the culture environment on the EV protein cargo (Supplementary Fig. 1A). This diversity suggests that the mechanical and structural cues inherent to each method shape the proteomic profile of EVs, which could have implications for their functional properties (Supplementary Fig. 1A). The number of shear stress-related proteins was quantified based on the literature reference list ^20,21^, showing the highest number in HS cultures, followed by 3D, and the least in 2D, suggesting that mechanical stress influences protein expression of shear stress-related proteins and cellular response to mechanical stimulus (Fig. 1C and Fig. 1D). Despite these differences, CD63, a key exosomal marker, exhibited similar expression levels across all methods, indicating consistency in certain aspects of exosome biology (Fig. 1E). In contrast, Piezo1 protein, associated with mechanotransduction, was significantly more expressed in HS compared to 3D, underscoring the impact of shear stress conditions on mechanosensitive protein expression (Fig. 1F and Supplementary figure 1B and 1C). These results were supported by the Western Blot analysis. Western blot analysis of tetraspanin EV markers (CD9, CD63, and CD81) showed indeed no noticeable difference between hMSC-EVs from 3D and HS cultures, suggesting that vesicle identity remains conserved (Fig. 1G). However, western blot results for Piezo1 protein confirmed its heightened expression in HS cultures (hMSC cell lysates) compared to 3D, further validating the role of shear stress in modulating mechanotransduction pathways (Fig. 1H). These findings collectively underscore the profound impact of culture conditions on EV production, particularly under shear stress.

**Figure 1.**
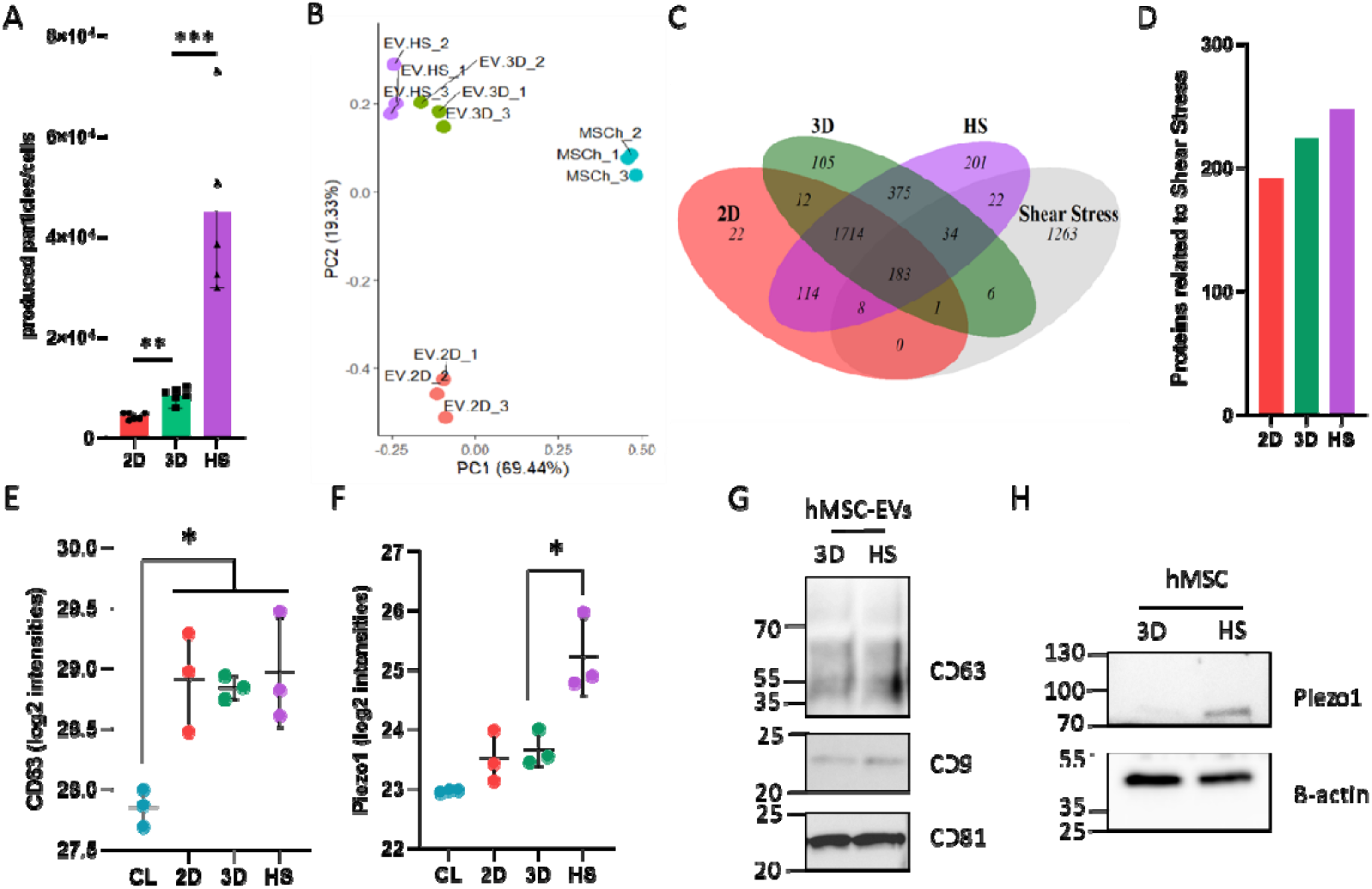
Production of EVs using different culture methods have distinct profiles. **(A)** Particle production per cell in conditioned media of 2D, 3D, and HS cultures of MSCs, six independent experiments were performed. Bars represent results from five independent replicates (*** p < 0.001, ** p < 0.01) **(B)** Principal component analysis (PCA) of proteomic data of MSCs cells and EVs isolated from MSCs in 2D, 3D and HS, three independent samples were analyzed. **(C)** Venn diagram depicting significantly expressed proteins unique or shear stress-related among the EVs isolated from MSCs in 2D, 3D and HS. **(D)** Number of shear stress-related proteins differentially expressed in EVs isolated from MSCs in 2D, 3D and HS. **(E)** Expression levels of CD63 in cell lysate (CL) of MSCs and EVs isolated from MSCs in 2D, 3D and HS. Bars represent results from three independent replicates (* p < 0.05) **(F)** Expression of Piezo1 protein in cell lysate (CL) of MSCs and EVs isolated from MSCs in 2D, 3D and HS. Bars represent results from three independent replicates (* p < 0.05) **(G)** Western blot analysis of tetraspanin EV markers (CD63, CD9, CD81) in 3D and HS cultures, with no observable differences between groups. **(H)** Western blot analysis of Piezo1 and b-actin protein expressions in MSCs.

### Activation of Piezo1 in 2D-cultured human MSC enhances the EV release while its inhibition does not alter EV production

To first access the impact of Piezo1 in enhancing EV secretion, human MSCs were cultured in 2D and treated with YODA1 and GsMTx4 (Figure 2). YODA1, a Piezo1 ion channel agonist, was selected through high-throughput screening of chemical libraries for compounds that modulate Piezo1 activity ^22^. YODA1 binds directly to Piezo1, stabilizing its open conformation and reducing the mechanical threshold required for channel activation ^22^. This allows the channel to remain active even under minimal mechanical stress, mimicking the effects of shear forces or tension on the plasma membrane ^23^. Treatment with YODA1 significantly increased dose dependent effect in particle production at both 50 µM and 100 µM concentrations, with the effect starting to be observed at 4 hours and maintained at 24 hours post-treatment. Importantly, the viability of the cells remained unaffected by these treatments, indicating that the increased particle production was not due to cytotoxicity (Fig. 2A). GsMTx4 (Grammostola mechanotoxin 4) was first isolated from the venom of the Chilean rose tarantula (*Grammostola rosea*) ^24^. It is a peptide toxin known for its ability to selectively inhibit mechanosensitive ion channels, including Piezo1. GsMTx4 functions by embedding itself into the lipid bilayer of the plasma membrane near mechanosensitive ion channels ^25^. This interaction alters the local membrane tension, preventing the channels from responding to mechanical stimuli ^26^. In contrast to YODA1, treatment with GsMTx4, did not alter EV production, suggesting that Piezo1 does not take part in the process of EV release in 2D condition when mechanical forces are not present. Similarly, cell viability remained stable following GsMTx4 treatment, further confirming that the experimental conditions did not compromise cellular health (Fig. 2B). These results highlight the role of Piezo1 activation in enhancing EV production without compromising cell viability.

**Figure 2.**
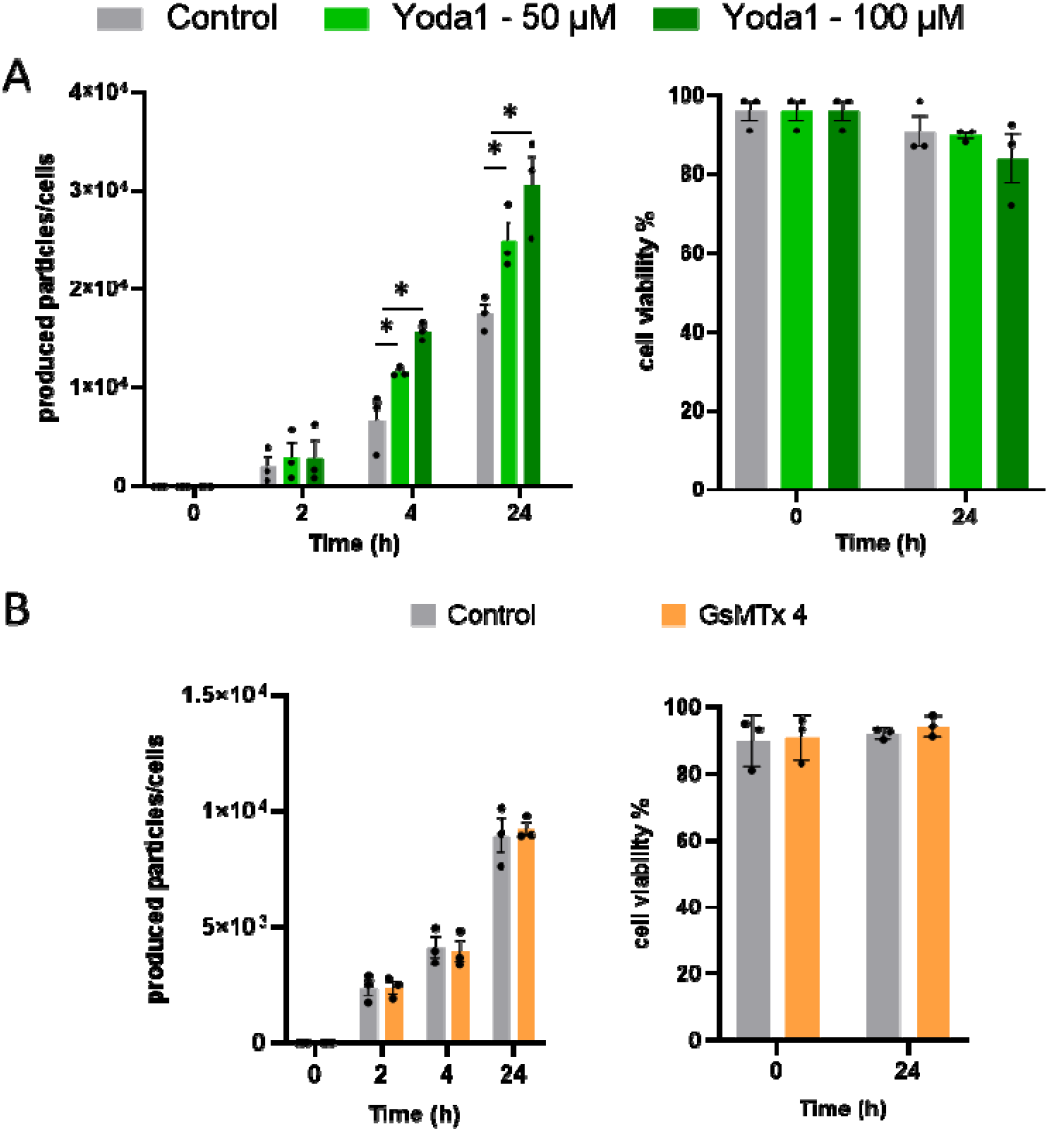
Modulation of Piezo1 activity impacts EV production in 2D-cultured human MSCs. (A) Particle production per cell and cell viability in 2D-cultured hMSCs after treatment with the Piezo1 agonist YODA1 at concentrations of 50 µM and 100 µM. (B) Particle production per cell and cell viability in 2D-cultured hMSCs after treatment with the Piezo1 inhibitor GsMTx4, demonstrating no change in EV production. Bars represent results from three independent replicates (* p < 0.05).

### Piezo1 is essential for the increase EV production by high shear stress in bioreactors

To explore how mechanical stimulation via Piezo1 activation influences the increased production of EVs, hMSCs were cultured in stirred-tank bioreactors to mimic specific mechanical stimulation. These bioreactors provided two distinct conditions: after cells grew on microcarriers, they were stirred on the one hand at minimal speed (3D), enough to maintain them in suspension, and on the other hand at high speed creating higher mechanical forces and high-shear (HS) conditions. These conditions allow for a closer examination of the role of Piezo1, a mechanosensitive ion channel, in sensing and responding to mechanical stimuli, thereby triggering cellular processes that enhance EV secretion. In 3D cultures, treatment with the Piezo1 inhibitor GsMTx4 did not affect EV production, suggesting that basal EV release is independent of Piezo1 activity under these conditions in a same way that in 2D cultures. However, treatment with YODA1, a Piezo1 agonist, significantly enhanced EV production also in a same manner as in 2D cultures, indicating that Piezo1 activation promotes vesicle release in 3D cultures (Fig. 3A). In HS cultures, with high-speed agitation, GsMTx4 inhibited EV production, demonstrating that shear-stress-induced EV release is mediated by Piezo1 activity. Consistently, treatment with YODA1 further increased EV production under 3D and HS conditions, highlighting an additive effect of Piezo1 activation on shear-induced vesicle release (Fig. 3B). This suggests that Piezo1 activity does not reach saturation under these conditions, as the EV production can still be further enhanced by the application of its agonist, YODA1. Comparative analysis of EV production, expressed as a percentage change relative to controls, confirmed these results. In 3D cultures, YODA1 significantly increased EV production and in HS cultures, YODA1 amplified EV release, whereas GsMTx4 effectively suppressed it (Fig. 3C). These results underscore the critical role of Piezo1 in mediating EV production in response to mechanical stimuli.

**Figure 3.**
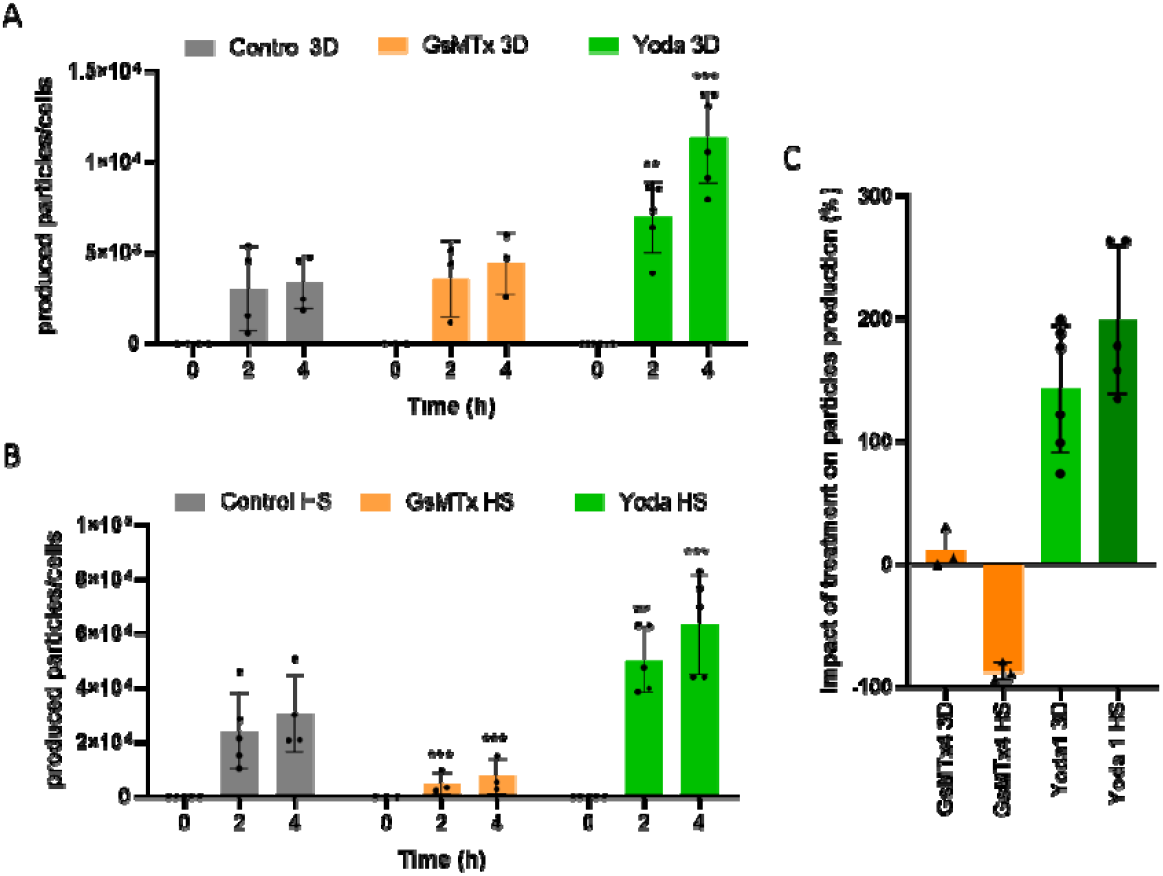
Piezo1 activity modulates EV production in 3D and high-shear (HS) bioreactor cultures. (A) Particle production per cell in MSCs cultured in a 3D stirred-tank bioreactor, showing no change with GsMTx4 (Piezo1 inhibitor) treatment but a significant increase with YODA1 (Piezo1 agonist). (B) Particle production per cell in MSCs cultured in HS conditions, where GsMTx4 treatment reduced EV production induced by shear stress, whereas YODA1 treatment further enhanced EV production. (C) Percentage change in EV production relative to controls, comparing the effects of GsMTx4 and YODA1 in 3D and HS cultures. For all graphs bars represent results from three or five independent replicates (*** p < 0.001, * p < 0.05).

MSC EVs have classical EV size and markers according to electron microscopy and nano-flow cytometry Electron microscopy images reveal the morphology of EVs produced by hMSCs cultured using 2D, 3D, and HS methods. The vesicles from all three culture conditions exhibit the characteristic round morphology and heterogeneous size distribution rof EVs. Despite differences in culture methods, the structural integrity and shape of the vesicles appear consistent across 2D, 3D, and HS samples, suggesting that the fundamental vesicle architecture is preserved regardless of the mechanical stimuli applied during culture. Due to lower concentrations, less vesicles were observed in the images from the control and treated with GsMTx4 conditions. No significant differences in the size of particles due to the treatments or method of culture was found (data not shown).

EVs across different culture methods were characterized regarding the expression of specific tetraspanins markers (CD9, CD63, and CD81) to detect the impact of Piezo1 activation and inhibition on their production. For CD9 expression, significant differences were observed in 3D cultures, where treatment with YODA1 resulted in higher expression compared to GsMTx4. However, no notable differences were observed in 2D or HS cultures for this marker (Fig. 4A). CD63 expression was influenced by Piezo1 modulation across all culture methods; YODA1 treatment increased expression compared to GsMTx4 in 2D, 3D, and HS cultures, indicating a consistent response to Piezo1 activation (Fig. 4B). In contrast, CD81 expression showed no significant differences between YODA1 and GsMTx4 treatments across any of the culture methods, suggesting that Piezo1 modulation does not impact this marker (Fig. 4C). These results highlight the differential effects of Piezo1 activity on specific tetraspanin markers under varying culture conditions.

**Figure 4.**
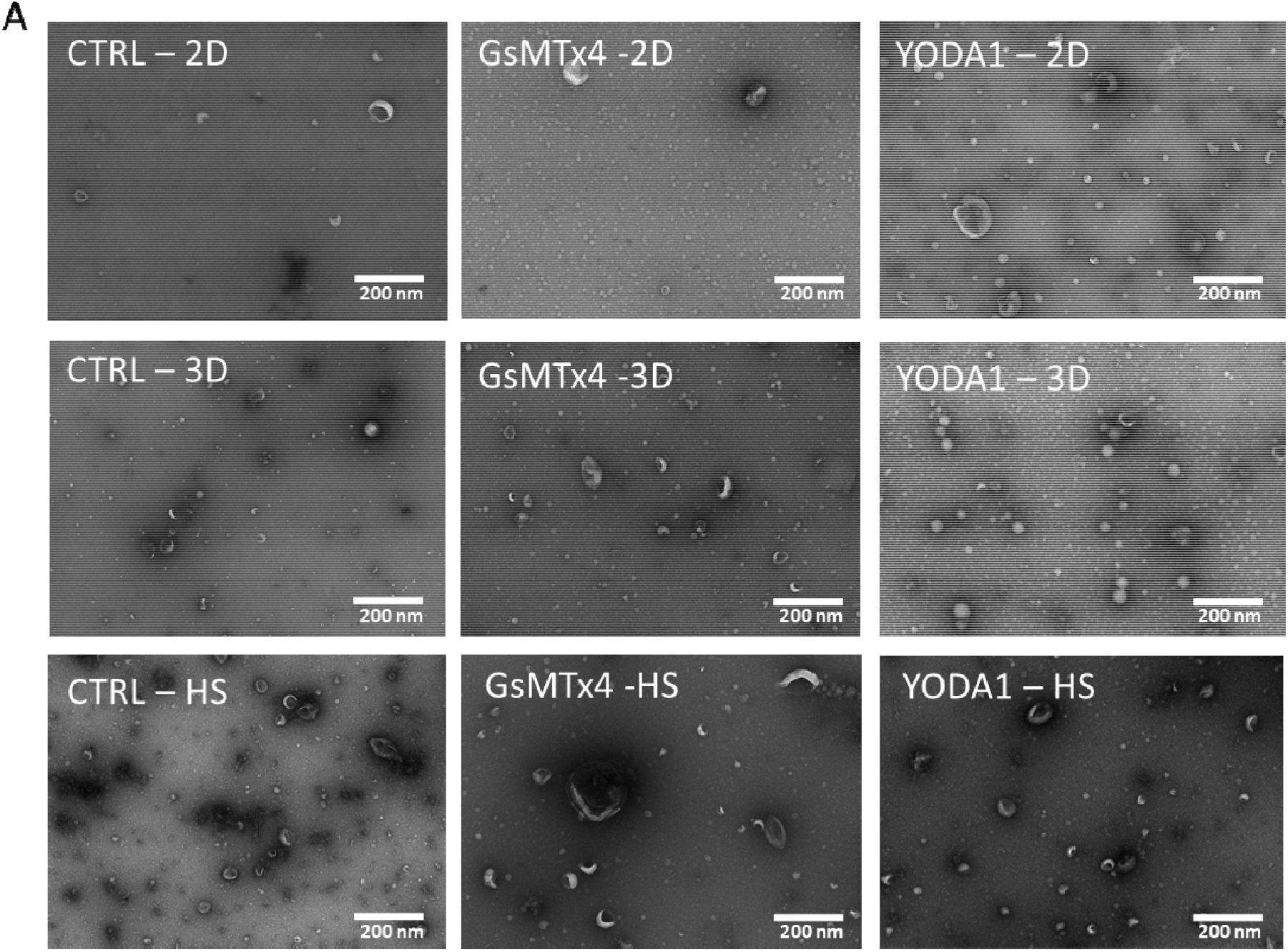
Morphology preservation of EVs produced under different culture conditions. Electron microscopy images of EVs derived from MSCs cultured in 2D, 3D, and HS conditions. Scale bars: 200 nm.

**Figure 4.**
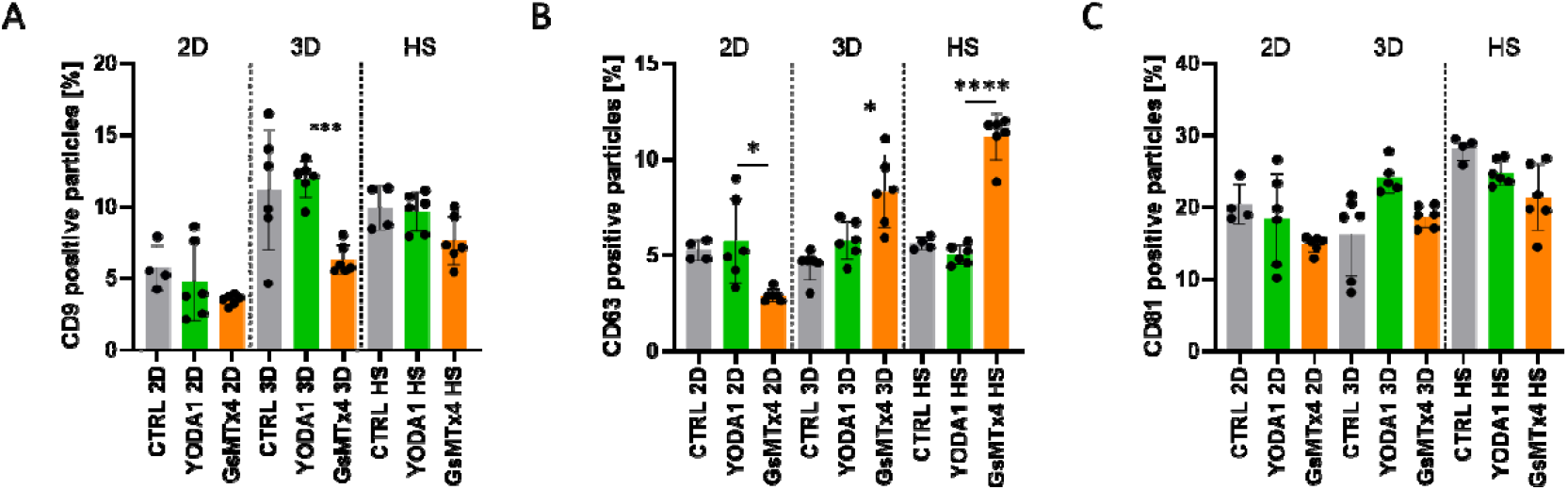
Impact of Piezo1 modulation on CD9 and CD63 expression across culture methods and treatments. (A) Percentage of CD9 expression in EVs from MSCs from 2D, 3D, and HS cultures. (B) Percentage of CD63 expression in EVs from MSCs. (C) Percentage of CD81 expression in EVs from MSCs. For all graphs bars represent results from three biological samples were analyzed in duplicate (**** p < 0.0001, *** p < 0.001, * p < 0.05).

## Discussion

This study highlights for the first time the critical role of Piezo1 ion channel activation in modulating EVs production after comparison across various culture methods, including 2D, 3D, and high-shear (HS) conditions. The results shown here demonstrate that both mechanical stimuli, such as shear stress, and chemical activation, using the Piezo1 agonist YODA1, significantly enhance EV secretion by human mesenchymal stem cells. Others also demonstrated that using shear stress stimulation and Piezo ion channel using YODA1 increases EV release by red blood cells ^27^. These findings collectively suggest that Piezo1 serves as a key mechanotransducer, linking external physical stimulation by a turbulent flow to the regulation of EV biogenesis.

Mechanotransduction is vital across many biological processes, including the development of neuronal cells, the sensation of pain, and the regulation of red blood cell volume^28,29^. When cells experience mechanical forces such as shear stress, the plasma membrane undergoes tension, which can activate ion channels and particularly Piezo channels in this context ^29^. They act as mechanosensors that open in response to increased membrane tension, allowing calcium ions to flow into the cell ^29^. This mechanical perturbation triggers a cascade of intracellular events, one of the most critical being the elevation of intracellular calcium ions (Ca^2+^) ^30^. Piezo1 and Piezo2 are mechanically activated ion channels integral to the process of mechanotransduction, where cells convert physical forces into biochemical signals ^31^. It is a large, trimeric transmembrane protein that forms a dome-shaped structure in the plasma membrane ^32^. When mechanical stimuli, such as shear stress, compression, or stretch are applied to the membrane, Piezo1 undergoes conformational changes that open the channel ^33,34^. This opening allows the influx of cations, primarily calcium ions (Ca^2+^), which act as secondary messengers to trigger downstream signaling cascades ^35^. As calcium levels rise, they play a pivotal role in facilitating lysosome exocytosis, a process where lysosomes fuse with the plasma membrane to release their contents, including EVs, into the extracellular space ^36^.

Here, we establish a correlation between Piezo1, a key regulator of mechanotransduction, and an increase of EV production, paving the way for advancements in EV-based technologies. Understanding and harnessing the interplay between mechanotransduction and EV biogenesis is a promising step toward realizing the full potential of EVs in regenerative medicine, drug delivery, and diagnostics. By leveraging mechanical and pharmacological stimuli, researchers can optimize EV production and tailor their functionality for specific therapeutic applications.

The morphological consistency of EVs across 2D, 3D, and HS culture conditions, as observed in electron microscopy images, supports the hypothesis that Piezo1 modulation does not compromise EV structure. However, Piezo1 activation appears to influence the expression of specific tetraspanin markers, such as CD9 and CD63, but not CD81, suggesting a selective role for Piezo1 in modulating EV surface protein composition. These observations could reflect differences in the pathways of EV biogenesis activated under varying culture conditions and stimuli. Further studies are needed to dissect these pathways and identify the molecular players involved in Piezo1-mediated EV production. Moreover, the significant impact of Piezo1 modulation in 3D and HS cultures suggests that biomechanical forces play a pivotal role in EV biogenesis, potentially mimicking physiological conditions more accurately than traditional 2D cultures.

EVs can be originated from the endosomal pathway and are formed as intraluminal vesicles (ILVs) within multivesicular bodies (MVBs)^37^. This step is mediated by the endosomal sorting complexes required for transport (ESCRT) machinery, lipid rafts, and tetraspanins. After their secretion, MVBs either fuse with lysosomes for degradation or with the plasma membrane to release ILVs as exosomes^38^. Mechanotransduction could influence exosome release by modulating intracellular calcium levels, which are essential for MVB trafficking and fusion with the plasma membrane. Also, due to elevated calcium levels, mechanotransduction could promotes cytoskeletal reorganization, a critical step in MVB transport to the plasma membrane, and facilitates vesicle secretion through SNARE complex activity^39^. It was already shown that increase of Ca^2+^ elevates EV release ^39,40^. Beyond Piezo1, other mechanosensitive pathways could influence EV biogenesis as integrins, which mediate cell adhesion and sense extracellular matrix stiffness ^29^ or as caveolae,membrane invaginations sensitive to membrane tension that may participate in EV formation under mechanical stress conditions ^41^.

These findings have significant implications for the field of regenerative medicine, drug delivery, and biomarker discovery, where EVs are increasingly being used as therapeutic tools and diagnostic biomarkers. Enhancing EV production through mechanical activation offers a scalable and efficient strategy to meet the growing demand for these vesicles. Additionally, understanding the role of Piezo1 in modulating EV composition opens avenues for a better comprehension of the mechanisms at play when cells undergo a mechanical stimulation via turbulent fluid. The growing recognition of EVs as powerful therapeutic agents underscores the importance of developing robust and scalable production methods and understanding how these processes may affect EV biogenesis. Advances in bioreactor technology, mechanotransduction research, and pharmacological modulation provide exciting opportunities to overcome existing barriers. Further research into optimizing EV production and understanding their underlying mechanisms will pave the way for their successful clinical translation.

The proposed mechanism for EV production through Piezo1 activation is summarized in a schematic representation. Mechanical stimuli, such as shear stress, activate the Piezo1 ion channel, leading to an increase in EV secretion. Similarly, chemical activation of Piezo1 using the agonist YODA1 also enhances EV production, mimicking the effects of mechanical stimuli. In contrast, blocking Piezo1 with the inhibitor GsMTx4 suppresses this mechanism, preventing the stimulation of EV secretion by either mechanical forces or YODA1 treatment (Fig. 5).

**Figure 5.**
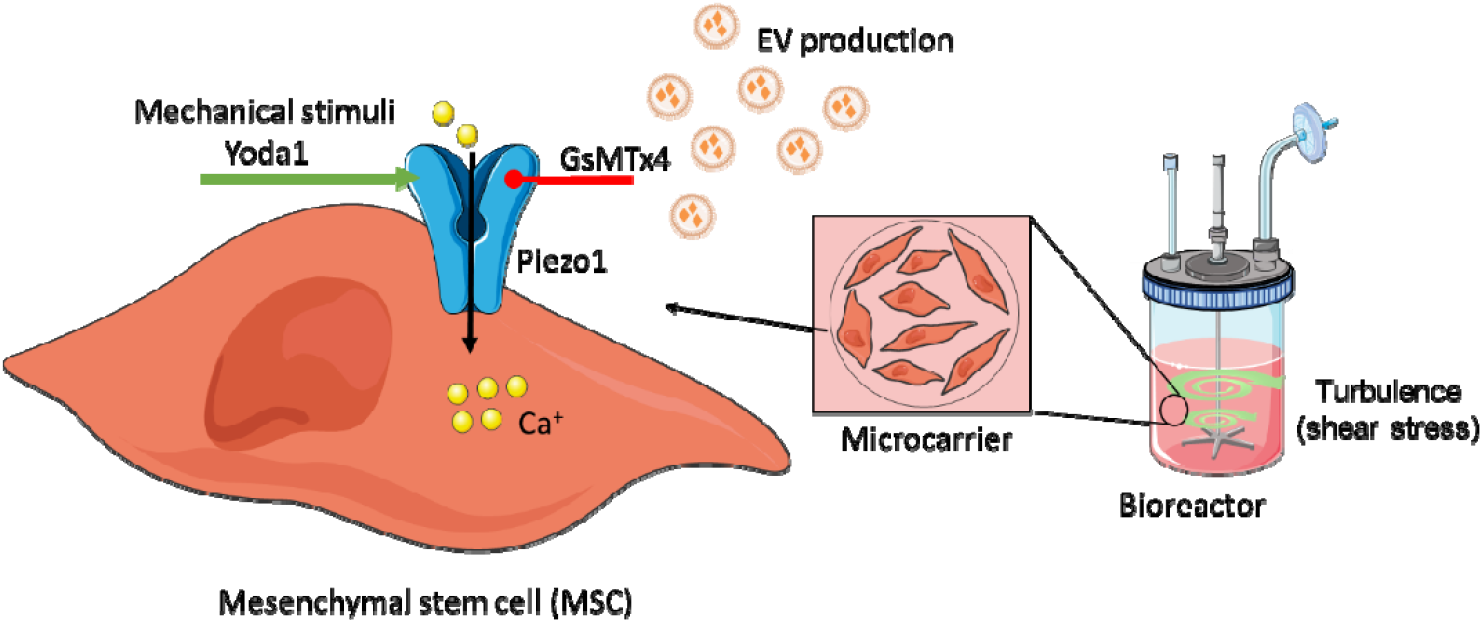
Proposed mechanism of EV production via Piezo1 activation. Schematic representation showing that mechanical stimuli or treatment with YODA1 (Piezo1 agonist) enhance EV secretion through Piezo1 activation. Blocking Piezo1 with GsMTx4 inhibits EV production induced by both mechanical and pharmacological activation.

In conclusion, this study establishes Piezo1 as a critical regulator of EV production under mechanical stimuli and provides a foundation for future research into the mechanotransduction pathways involved in EV biogenesis. By leveraging mechanical and chemical stimuli, it may be possible to optimize EV production and functionality, addressing key challenges in the field and unlocking the full potential of EV-based technologies.

## Material and methods

### Cell culture and EV production

Human adipose tissue-derived stem cells (Cell-Easy, France) were expanded in culture flasks with complete media (aMEM supplemented with 10% Bovine Fetal Serum (FBS)) and kept at 37°C in incubators. Microcarriers (200 µm, Cytodex 1, Cytiva, USA) were sterilized by autoclaving after rehydration in Phosphate Buffered Saline (PBS). Before cell seeding, microcarriers were incubated in complete aMEM at 37°C. Cells were seeded onto the microcarriers at a ratio of 5:1 and subjected to 24 cycles of 45 minutes of rest and 5 minutes of gentle mixing at 100 Lk in a Dasbox bioreactor (Eppendorf, Germany). After intermittent cycles, the cells were submitted to continuous agitation at 100 Lk until they reached confluence. Once confluent, cells on microcarriers were washed five times with PBS and resuspended in serum-free aMEM before EV production was initiated. The impeller speed was increased to 34 Lk for 4 hours, generating shear stress for EV production. For mass spectrometry analysis, spinner flasks were used to produce EVs. A comparison of the production in spinner flasks or in stirred-tank bioreactors showed no impact of the culture system on EV production (Supplementary figure 2). The conditioned media (CM) was sampled at the start of the shear stress stimulation and after 2 and 4 hours. EV production was also performed by adding YODA1 or GsMTx4 (Merk, USA) For the clearance of CM, cells were firstly removed with a cell strainer and cellular debris were then removed by centrifuging the supernatant at 3000 g for 15 minutes.

To evaluate the turbulence in the applied flow, we calculated the forces acting in this phenomenon to make it transferable among different culture systems (eg. Spinner flasks and stirred tank bioreactors and different culture volumes). Turbulence occurs when the Reynolds number (Re) exceeds 10,000, calculated as *Re* = *vL/V*, where v is velocity, L is flow size, and V is kinematic viscosity. In turbulent flow, energy cascades from large to small eddies until viscous forces dissipate. The smallest eddy size, or Kolmogorov length (Lk), is determined by 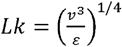, where *ε* is energy dissipation, estimated as *ε* = *PoN*^*3*^*D*^*5*^*/V*, where Po is the power number, D is the impeller diameter, N is the speed in rpm and V is the volume. Viscosity (v) considers media and bead properties, and bead swelling determines volumic fraction (Φ\Phi). Shear stress on microcarrier surfaces is calculated based on Lk and microcarrier diameter. These calculations were adjusted, as according to impeller speed and geometry, to optimize turbulence for efficient EV production.

### Cell viability

To quantify dead and live cells in 2D cultures the Nucleocounter NC-200 (Chemometec, Denmark) device was used according to manufacturer with calibrated cassettes from the supplier containing Acridin Orange and DAPI.

### EV isolation

For EV enrichment and purification, a total CM of 50 or 100 mL from one production was thawed. This CM was then concentrated 50 or 100 times using a 100 kDa cut-off hollow fiber modified polyethersulfone (mPES) membrane filter column (Repligen, USA) operated manually. The permeability of the hollow fiber was always checked previously to the purification using a PALL TFF system (PALL Corporation, USA). For buffer exchange and removal of proteins, the concentrated EVs were diafiltered in the system with 5-times their volume of PBS.

### Characterization of EVs

EV suspensions were examined using a ZetaView (Particle Metrix, Germany) according to the manufacturer’s instructions to evaluate their concentration, size distribution, mean and mode. Samples were diluted according to the detection limit of the instrument with 0,22 µm filtered PBS. The instrument uses the nanoparticle tracking analysis (NTA) method to link the Brownian motion of EVs to their size. NTA post-acquisition settings were optimized and kept consistent for repeated analyses.

### Nano-flow cytometry

Nano-flow cytometry was conducted using the Flow NanoAnalyzer (NanoFCM, UK) according to the manufacturer’s instructions. Calibration involved using 200 nm PS QC beads (NanoFCM, UK) for particle concentration and S16M-Exo silica beads (NanoFCM, UK) to create a calibration curve for converting side scatter intensities to particle sizes. The laser setup included 488 nm and 640 nm lasers at 20 mW with lens filters at 525/40 and 670/30. For staining, Alexa Fluor® 647-conjugated monoclonal mouse antibodies targeting CD63, CD9, and CD81 (Santa Cruz Biotechnology, USA) were used, along with isotype controls. Antibodies were prepared by centrifugation at 10,000g for 15 minutes or filtration through a 0.22 µm syringe filter. Staining was performed overnight at 4°C with a ratio of 1:10.. Before measurement, labeled vesicles were diluted 1:50 or 1:20 times. Positivity was defined using PBS alone, unstained vesicles, isotype controls, and auto-thresholding. Initial sample acquisition determined particle concentration, which was then adjusted to 5.10^9^ particles/mL for optimal EV staining. Blank controls included PBS and antibodies diluted 100-fold in PBS. To set the threshold between positive events and background noise in the fluorescence channel, signals from unstained EVs and EVs with isotype control were used. Each measurement was performed in triplicate, and data were analyzed using NF Profession 1.0 software.

### Western blot

The protein concentration of CM and concentrated EV samples was measured using the micro BCA (Invitrogen, USA) following the manufacturer’s instructions. Optical density (OD) at 562 nm was read on plate reader. For SDS-PAGE analysis, samples were prepared with 4X Laemmli buffer (Bio-Rad, USA) containing 50 µM dithiothreitol (DTT) as a reducing agent. Samples were run on 4–20% SDS-PAGE gels (Bio-Rad, USA) using a Protean Mini System (Bio-Rad, USA). Molecular weight was estimated using Protein Standard (Bio-Rad, USA). Proteins were transferred to polyvinylidene difluoride (PVDF) membranes using the Trans-Blot Turbo Transfer System (Bio-Rad, USA). Membranes were blocked with tris-buffered saline (TBS) containing 0.05% Tween 20 and 2% bovine serum albumin (BSA). Specific proteins were detected using monoclonal antibodies: CD63, CD81, CD9 (Cell Signaling, USA), Piezo1 and b-actin (Proteintech, USA). Detection involved HRP-conjugated secondary antibodies: rabbit anti-mouse IgG (Abcam, UK). Specific bands were visualized and quantified using Clarity Western ECL substrate (Bio-Rad, USA).

### Transmission electron microscopy (TEM)

To visualize EVs in TEM, samples were submitted to negative staining protocol. 4□µL of isolated EV samples were added on copper grids F/C (300 mesh) and incubated at room temperature for 5□min. The excess was removed with a filter paper before staining with 1% uranyl acetate for 20□s at room temperature. After removal of excess uranyl acetate, the samples were dried and visualized using a HT 7700 120kV (Hitachi, Japan).

### NanoLC-MS/MS protein identification and quantification

S-TrapTM micro spin column (Protifi, Hutington, USA) digestion was performed on EV and cell lysate from ASCh cell line according to manufacturer’s protocol. Briefly, 5% SDS was added to the samples. Proteins were alkylated with the addition of chloroacetamide to a final concentration of 50mM. Aqueous phosphoric acid was added to a final concentration of 1.2%. Colloidal protein particulate was formed with the addition of 6 times the sample volume of S-Trap binding buffer (90% aqueous methanol, 100mM TEAB, pH7.1). The mixtures were put on the S-Trap micro 1.7mL columns and centrifuged at 4,000g for 30 seconds. The columns were washed four times with 150µL S-Trap binding buffer and centrifuged at 4,000g for 30 seconds with 180 degrees rotation of the columns between washes. Samples were digested with 0.8 µg of trypsin (Promega) at 47°C for 1h30.

Samples were resuspended in 20 µL of 2% ACN, 0.1% FA in HPLC-grade water. Each sample was injected three times. For each run, 1 µL was injected in a nanoRSLC-Q Exactive PLUS (RSLC Ultimate 3000) (Thermo Scientific,Waltham MA, USA). Peptides were loaded onto a µ-precolumn (Acclaim PepMap 100 C18, cartridge, 300 µm i.d.×5 mm, 5 µm) (Thermo Scientific), and were separated on a 50 cm reversed-phase liquid chromatographic column (0.075 mm ID, Acclaim PepMap 100, C18, 2 µm) (Thermo Scientific). Chromatography solvents were (A) 0.1% formic acid in water, and (B) 80% acetonitrile, 0.08% formic acid. Peptides were eluted from the column with the following gradient 5% to 40% B (120 minutes), 40% to 80% (1 minutes). At 121 minutes, the gradient stayed at 80% for 5 minutes and, at 126 minutes, it returned to 5% to re-equilibrate the column for 20 minutes before the next injection. One blank were run between each replicates to prevent sample carryover. Peptides eluting from the column were analyzed by data dependent MS/MS, using top-10 acquisition method. Peptides were fragmented using higher-energy collisional dissociation (HCD). Briefly, the instrument settings were as follows: resolution was set to 70,000 for MS scans and 17,500 for the data dependent MS/MS scans in order to increase speed. The MS AGC target was set to 3.106 counts with maximum injection time set to 200 ms, while MS/MS AGC target was set to 1.105 with maximum injection time set to 120 ms. The MS scan range was from 400 to 2000 m/z. Dynamic exclusion was set to 30 seconds duration.

### Data analysis

The obtained data were analyzed using MaxQuant version 2.0.1.0 and searched with Andromeda search engine against the UniProtKB/Swiss-Prot Homo sapiens database (release 02-04-2020, 20365 entries). To search parent mass and fragment ions, we set a mass deviation of 3 ppm and 20 ppm respectively. The minimum peptide length was set to 7 amino acids and strict specificity for trypsin cleavage was required, allowing up to two missed cleavage sites. Carbamidomethylation (Cys) was set as fixed modification, whereas oxidation (Met) and N-term acetylation were set as variable modifications. The false discovery rates (FDRs) at the protein and peptide level were set to 1%. The reverse and common contaminants hits were removed from MaxQuant output. Proteins were quantified according to the MaxQuant label-free algorithm using LFQ intensities; protein quantification was obtained using at least 2 peptides per protein. The mass spectrometry proteomics data have been deposited to the ProteomeXchange Consortium via the PRIDE ^42^ partner repository with the dataset identifier PXD059823.

The protein intensity values (log2) were normalized using the median absolute deviation (MAD) centering approach (Supplementary figure 3A). The consistency among biological replicates was assessed by calculating Pearson correlation coefficients between log2-transformed matrices (Supplementary figure 3B). Principal component analysis was conducted to explore variance patterns across sample phenotypes after normalization. Missing data were imputed utilizing the DEP package in R ^43^. Differential protein expression analysis was performed through linear modeling of the normalized dataset using the limma package ^44^. Proteins were considered significantly differentially expressed if they had an adjusted p-value of less than 0.05 (FDR). The analysis was executed in R software (version 4.2.3) within the RStudio environment, enabling visualization of differential proteins through heatmaps, hierarchical clustering, and volcano plots, as detailed in (https://github.com/41ison).

## Acknowledgments

AKAS has received funding from the European Research Council (ERC) under the European Union Horizon 2020 Research and Innovation Program (grant agreement no. 852791).

This work was possible thanks to the IVETh facility supported by the IdEx Université Paris Cité, ANR-18-IDEX-0001, by the Region Ile de France under the convention SESAME 2019 – IVETh (EX047011) and via the DIM BioConvS, by the Région Ile de France and Banque pour l’Investissement (BPI) under the convention Accompagnement et transformation des filières projet de recherche et développement N° DOS0154423/00 & DOS0154424/00, DOS0154426/00 & DOS0154427/00, and Agence Nationale de la Recherche through the program France 2030 “Integrateur biotherapie-bioproduction” (ANR-22-AIBB-0002). This project was also supported by fundings from CNRS and from the European Research Council (ERC) under the European Union’s Horizon 2020 research and innovation program (grant agreement No. 852791). We thank Christine Longin from electronic microscopy platform MIMA2. Also we thank for Alison F. A. Chaves in assisting in proteomic data analysis.

