## Supplementary figures for "Piezo ion channel activation increases the release of therapeutic extracellular vesicles after mechanical stimulation in bioreactors"

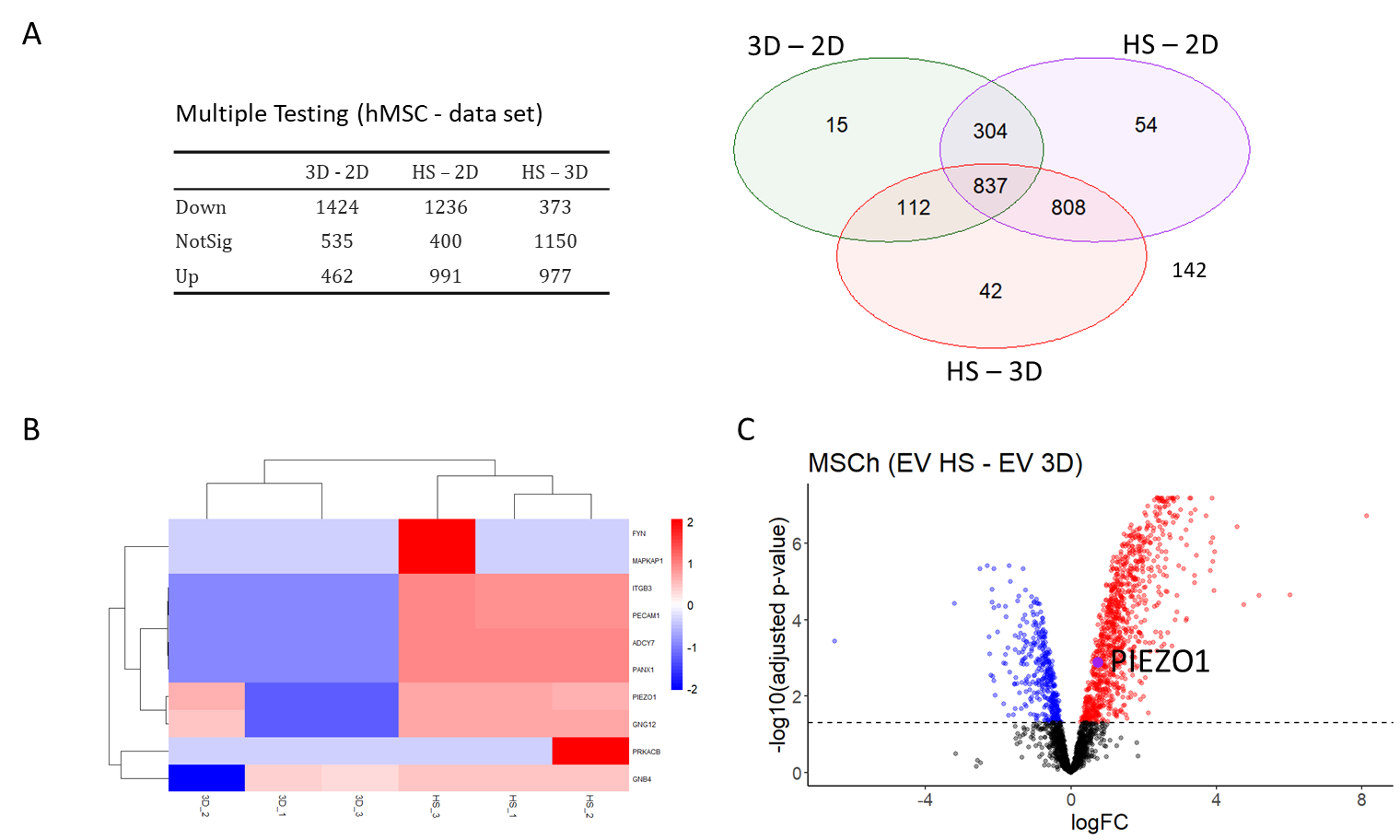

**Supplementary figure 1.** Differential Expression of Proteins Related to Cellular Response to Mechanical Stimulus in EV 3D vs. EV HS. (A) Venn diagram depicting significantly expressed proteins unique to or shared among the EVs isolated from MSCs in 2D, 3D and HS. (B) Heatmap displaying differentially expressed proteins associated with the cellular response to mechanical stimulus from Reactome database (R-HSA-2262752) in extracellular vesicles (EV) derived from 3D culture (EV 3D) compared to heat shock-treated EVs (EV HS). Rows represent proteins, and columns represent biological replicates. Color intensity corresponds to expression levels, with red indicating upregulation and blue indicating downregulation. (C) Volcano plot illustrating the differential expression of proteins in EV 3D versus EV HS. The x-axis represents the log2 fold change, and the y-axis represents the -log10(p-value). Significantly upregulated proteins are highlighted in red, while downregulated proteins are in blue. PIEZO1, a mechanosensitive ion channel, is prominently overexpressed in EV 3D and is specifically highlighted in the plot.

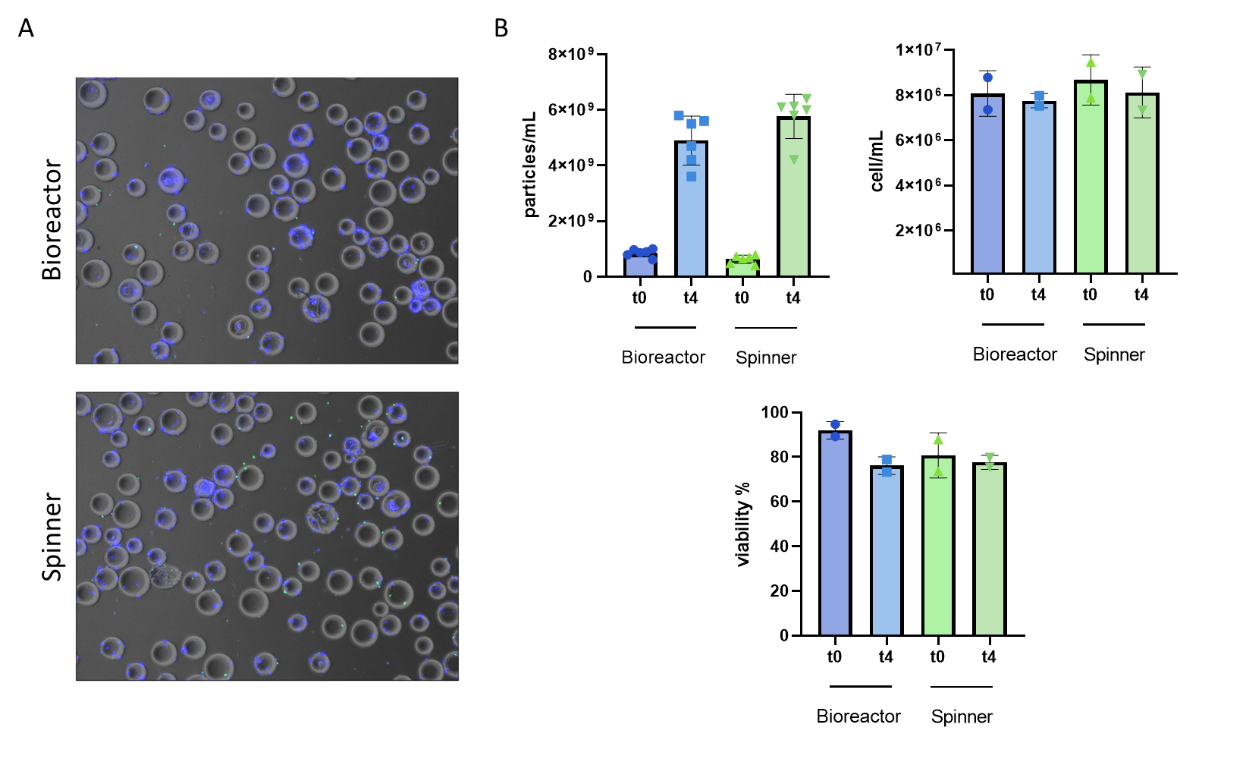

**Supplementary Figure 2.** MSCs on Microcarriers in Different Culture Systems. (A) Representative fluorescence microscopy images of mesenchymal stem cells (MSCs) cultured on microcarriers in a spinner flask (left) and a stirred-tank bioreactor (right). Nuclei are stained blue with DAPI, and activated Caspase-3, indicating apoptosis, is shown in green. (B) Quantitative analysis of particles per mL, total cell number, and cell viability at 0 and 4 hours of culture. Data are presented as mean ± standard deviation of three independent experiments.

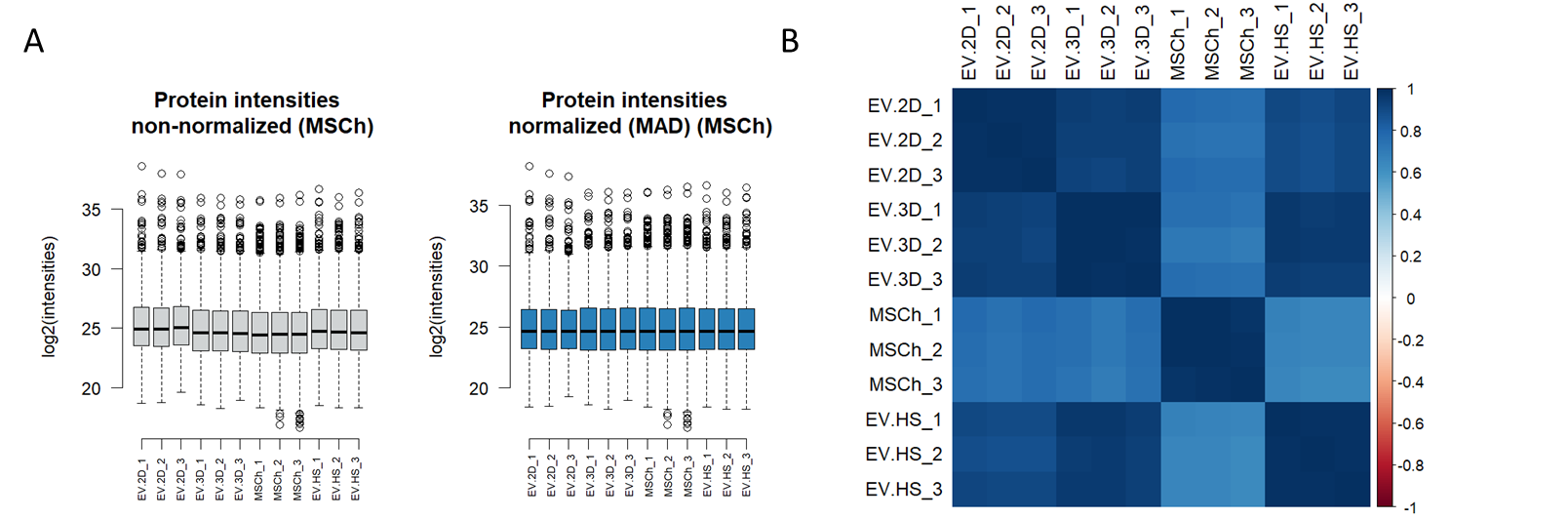

**Supplementary Figure 3.** Protein Intensity Normalization and Sparsity Assessment. (A) Protein intensity distributions before and after normalization. Raw intensities (left) show variability across samples, while normalized intensities (right) demonstrate improved consistency, ensuring comparability across conditions. (B) Reproducibility assessment of normalized protein intensities using Pearson correlation coefficients. The correlation matrix illustrates the consistency between biological replicates.
